# Outer membrane β-barrel structure prediction through the lens of AlphaFold2

**DOI:** 10.1101/2022.10.09.511469

**Authors:** Annika Topitsch, Torsten Schwede, Joana Pereira

## Abstract

Most proteins found in the outer membrane of Gram-negative bacteria share a common domain: the transmembrane β-barrel. These outer membrane β-barrels (OMBBs) occur in multiple sizes, and different families with a wide range of functions evolved independently by amplification from a pool of homologous ancestral ββ-hairpins. This is part of the reason why predicting their three-dimensional (3D) structure, especially by homology modeling, is a major challenge. Recently, DeepMind’s AlphaFold v2 (AF2) became the first structure prediction method to reach close-to-experimental atomic accuracy in CASP even for difficult targets. However, membrane proteins, especially OMBBs, were not abundant during its training, raising the question of how accurate the predictions are for these families. In this study, we assessed the performance of AF2 in the prediction of OMBBs of various topologies using an in-house-developed tool for the analysis of OMBB 3D structures, *barrOs*. In agreement with previous studies on other membrane protein classes, our results indicate that AF2 predicts OMBB structures at high accuracy independently of the use of templates, even for novel topologies absent from the training set. These results provide confidence on the models generated by AF2 and open the door to the structural elucidation of novel OMBB topologies identified in high-throughput OMBB annotation studies.

## Introduction

Protein structure prediction is an important tool to gain insights into the function and biological role of macromolecular machines from three-dimensional (3D) models. While the number of known natural protein sequences has been increasing exponentially [1,2] since the first sequencing of a protein in the 1950s, the experimental determination of macromolecular 3D structures is a laborious task. For this reason, and even with the recent considerable improvements in experimental biophysical methods, the rate by which protein structures are deposited in the Protein Data Bank (PDB) [3] is much lower than that by which protein-coding sequences are made available through GenBank or the UniProt Knowledgebase. One way of tightening this gap is to use computational approaches such as homology modeling, threading, or *ab initio* methods for protein structure prediction [4–7].

The Critical Assessment of Protein Structure Prediction (CASP) [8] experiment provides a platform for the benchmarking of such methods and, since its onset in the early 1990s, has fostered the development of multiple approaches exploring a wide range of data sources and computational techniques. Until recently, homology modeling was the method of choice to model 3D structures of proteins with homologs of known structure in the PDB, while *ab initio* methods were preferred for all others. However, *ab initio* modeling was rarely able to reach for such difficult targets the same level of accuracy as that typically reached by homology modeling. That changed in 2020, with DeepMind’s second version of the AlphaFold algorithm (AF2) providing, on average, close to experimental accuracy levels for most targets in the 14^th^ round of CASP (CASP14) [9,10], and later significantly expanding the structural coverage of the cataloged protein sequence space [11,12].

AF2 is a deep neural network with two attention-based transformation modules, where evolutionary, physical, and geometric information is used to perform end-to-end protein structure prediction [10]. The first module, the Evoformer, uses information from multiple sequence alignments (MSAs) and templates to generate a pair representation, a contact map of sorts, for the input. The second module, the structure module, uses this representation and the input sequence to fold the target protein. The network has been trained with all protein structures in the PDB as of April 30, 2018, and it is not tailored to any specific class of proteins.

However, the PDB is biased towards those proteins that are ‘easier’ to experimentally characterize, with only 10 % of its content corresponding to membrane proteins [13]. For this reason, it is not expected that AF2 is able to predict the 3D structure of transmembrane proteins as accurately as soluble ones, especially when multiple domains are present. In a recent study, Hegedűs *et al.* assessed AF2 structure prediction of α-helical transmembrane proteins. They observed that the models predicted by AF2 exhibited a known fold of α-helical transmembrane proteins for all 1,137 test cases, suggesting that the prediction of transmembrane proteins by AF2 is as accurate as for soluble proteins [14].

In this short report, we focus on the second-largest class of transmembrane proteins: the outer membrane β-barrels (OMBBs). OMBBs are abundant in Gram-negative bacteria, but are also found in chloroplasts, mitochondria and mitochondria-associated organelles [15–17]. They have both medical and biotechnological importance [18–20] as they are composed of an antiparallel β-sheet that connects back to itself to form a pore that crosses the outer membrane, where they perform a large variety of biological activities essential for survival [21]. They are found in a large spectrum of protein families either as single domains, together with other domains, or in multiple copies in the same chain [22]. Different OMBB families are composed of different numbers of β-strands [22,48]. The diameter of the barrel depends on the numbers of β-strands, but also on the shear number, which is, simply put, a measure of the parallel displacement of the strands relative to each other [23–25].

OMBB structure prediction is a challenging task as they can be traced back to a pool of homologous ancestral ββ-hairpins and novel families emerge by the reuse and amplification of smaller pieces from other OMBBs [26–29]. In the special case of homology modeling, when dealing with a novel family for which no full-length template is known or for which the full-length template corresponds to part of a larger β-barrel, the resulting model will either correspond to (1) a mix of local matches with mismatching shears that prevent the proper closing of the barrel, or (2) an incomplete, open barrel incompatible with the membrane environment. Current OMBB modeling approaches circumvent these problems by using external information specific to these proteins. These include the generation of perfect barrel structures directly from a theoretical description of a barrel [15,23,25,30], the prediction of transmembrane segments from sequence features [31–36] and their fitting into a putative membrane [37], and the prediction of contacts between those segments from free energy potentials based on statistical models [38–41] or evolutionary couplings [42,43].

In this short report, we sought to evaluate how the family-agnostic AF2 network performs for OMBBs. As of the time of AF2 training, about 100 single-chained OMBBs at a maximum of 70 % sequence identity (table S1) and with 8 up to 26 β-strands were deposited in the PDB. In the meantime, the structure of a 36-stranded OMBB, the translocon of the Fibrobacteres-Chlorobi-Bacteroidetes type 9 secretion system, was solved by cryo-EM [44], and more than 30 previously unknown OMBB families were predicted at the sequence level, including the largest ever reported OMBB, with at least 38 predicted strands [22]. In the case of long-known OMBB topologies, structural information has been fed into the network during the training phase. Thus, even without using homologous structures for modeling, OMBB models of high accuracy are expected. But since the newest topology of a 36-stranded OMBB was discovered after the date limit for inclusion in the training set, it is unclear how AF2 performs in such cases and what the impact of using templates is.

## Methods

### Collection of test case structures

Ten OMBBs of known structure, covering topologies of 8- to 24-stranded barrels (table S1) were first used as input for searches against an HHM database of the PDB70 (as of February 2021) with HHpred [45] through the MPI Bioinformatics toolkit [46]. Default parameters were used and all PDB chains matched at a p-value better than 10 collected. These were then analyzed with *barrOs* in order to (1) identify the matched PDB IDs that carry a barrel fold, (2) extract geometric features of the barrel region, and (3) extract the sequence of the barrel domains. With this, 129 unique OMBBs of known structure were collected and the sequences of the detected barrel regions, which include the barrel-forming strands and the connecting loops, were used as input to AF2.

### Identification of OMBB folds and extraction of barrel geometric features with *barrOs*

*barrOs* (for *barrel circle searcher*) is an in-house-developed tool that, given a PDB structure, uses a graph-based approach to identify the strands that form a barrel fold and then uses them to compute geometric features. This includes the number of strands, the barrel diameter, and the shear number of the barrel region. The method is family-agnostic and can take as input (1) a PDB structure, (2) a list of PDB IDs, or (3) HHsearch output files. It can be targeted specifically to transmembrane proteins by using the Orientations of Proteins in Membranes (OPM) database [47] as a source of 3D structures, and to OMBBs specifically by combining it with the results from HHsearch.

For each structure to be analyzed, *barrOs* starts by extracting all Cα atoms and searches for all β-strands. This is done by (1) running DSSP [48,49] and, in parallel, (2) detecting what we denote as ‘regular regions’. Regular regions are continuous backbone segments where the angle between the Cαi-Cαi+2 and Cαi+1-Cαi+3 vectors is lower than 25°. Regular and stranded region annotations derived from the DSSP (‘E’) output are then fused, and the resulting continuous intervals referred to as ‘strands’.

Two strands where the minimum interstrand distance of their Cα atoms is less than 5 Å are considered adjacent, allowing the construction of a strand-connectivity matrix. This matrix is then used to build an undirected, labeled graph, and the *cycle_basis* function implemented in *NetworkX* [50] is used to identify the nodes, i.e., the strands, that form a closed cycle. Given that OMBBs typically have an even number of strands (except for the 19-stranded mitochondria-specific OMBBs), if the resulting number of barrel-forming strands (the estimated topology) is uneven, this process is repeated using the regular regions or the DSSP-extracted strands until an even topology is obtained. If the result remains uneven, that topology is considered. Structures without detected structured barrels are excluded, and only those with a detected barrel fold are used for further analysis. This includes the estimation of the barrel height, average diameter and shear number, as defined in Murzin *et al.* [23].

### Running AF2

AF2 models were generated for the 129 OMBBs in three independent experiments with AlphaFold v2.0.1 on a local cluster instance with three different parameter settings: The first was performed with the default pipeline, which includes the use of all templates found in the PDB (labeled ‘M’). In the second, AF2 was run without considering any templates by setting the `--max_template_date` option to 1900-01-01 (labeled ‘Mnotemp’). And in the third, template usage was partially allowed by setting the `-- max_template_date` option to one day prior to the respective release date in order to exclude the deposited structure from being used as a template for modeling (labeled ‘Mreldate’).

### Model comparison and visualization

All AF2 models were used as input files for *barrOs* to identify barrel topologies and extract geometrical information of the barrel domains. Calculations of the template modeling score (TM-score) and the root-mean-square deviation (RMSD) were carried out with TM-align [51]. OpenStructure was used to calculate the per-residue local distance difference test score (Cα-lDDT) for each model [52].

## Results

As a first step, we evaluated how well AF2 captures the core geometric features of OMBBs in the presence and absence of templates, especially the topology of the domain, the average diameter of the channel and the shear, which measures the extent by which the strands are staggered (fig. 1). For that, the 129 experimental structures and the corresponding AF2 models were used as input for *barrOs.* The first observation is that AF2 predicted models with the correct topology for most cases; out of the 129, there were only two test cases where the number of strands in the model deviated by ±1. In these cases, visual inspection highlighted that the difference is not a result of an incorrectly modeled topology, but due to minor differences in the experimental structure and the AF2 model that misled *barrOs* during the identification of regular regions (figs. S1A-B).

**Figure 1.**
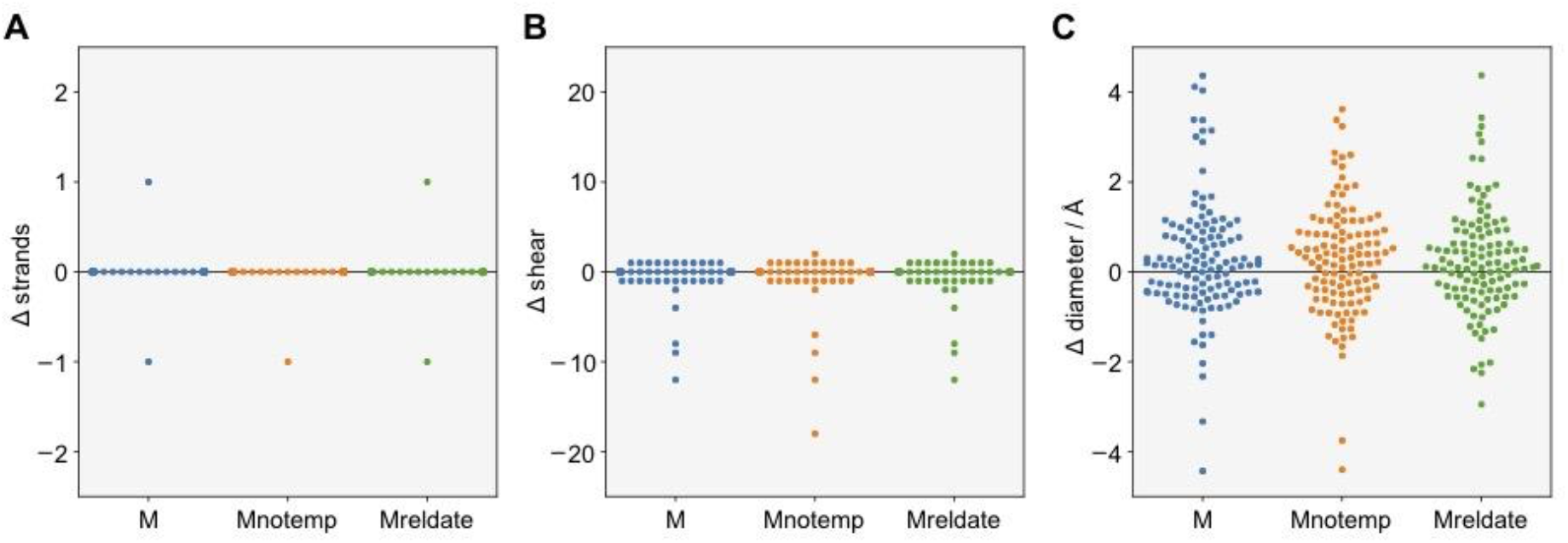
Predicted OMBB topologies and geometries. Differences (Δ) of the number of strands (A), shear numbers (B) and barrel diameters (C) in target structures and AF2 models. Data of AF2 models generated with using templates (‘M’), without using templates (‘Mnotemp’) and with using templates up to the release date of the target structure (‘Mreldate’), is shown in blue, orange and green, respectively.

Regarding the shear and barrel diameter, there are also only marginal differences between the target structures and the models predicted by AF2, with some noteworthy exceptions. One striking case is that of *Vibirio cholerae* OmpT (PDB ID 6EHD), where the shear of the model generated with no templates (‘Mnotemp’) was 18 residues smaller (fig. S1C). The reason here lies on an extracellular loop that in both AF2 models predicted using templates (‘M’ and ‘Mreldate’) and the experimental structure is modeled inwards, facing the channel of the barrel, while in the ‘Mnotemp’ model it faces the exterior, extending the strands that build the barrel region and leading to an incorrect value of the shear.

The agreement between the geometric features of AF2 models and their target experimental structure is also corroborated by superposition-based and superposition-free quality metrics. In the case of superposition-based metrics, high median TM-scores, and correspondingly low RMSD values, were observed for all three experiments (fig. 2A-B). The highest median TM-scores (0.98 ± 0.02) were obtained with the AF2 default pipeline, in which template information is used for the prediction of models (‘M’), but also when templates up to the release date (‘Mreldate’) were allowed. Excluding templates completely (‘Mnotemp’) only lead to marginally, and not statistically significant, lower TM-scores (0.97 ± 0.02).

**Figure 2.**
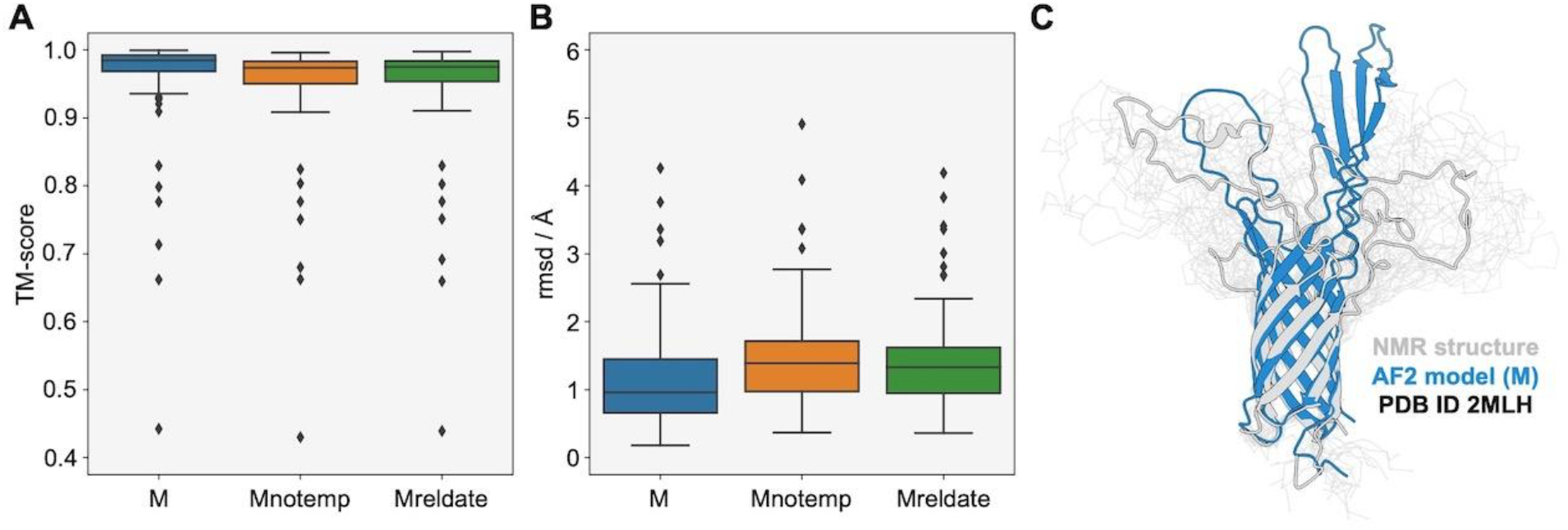
Full-length assessment of target structures and AF2 models. (A) The median TM-scores of the ‘M’, ‘Mnotemp’ and ‘Mreldate’ experiments are 0.98 ± 0.02, 0.97 ± 0.02 and 0.98 ± 0.02, respectively. (B) The median RMSD values, as computed by TMalign, of the ‘M’, ‘Mnotemp’ and ‘Mreldate’ experiments are 1.0 ± 0.5, 1.4 ± 0.5 and 1.3 ± 0.5 Å, respectively. (C) Solution NMR structure and AF2 model of PDB ID 2MLH shown in gray and blue, respectively. The backbone traces of the other 19 calculated conformers are shown in light gray.

This testifies to an overall high accuracy of the AF2 models independently on the use of templates, yet there are a few outliers below and above the lower and upper quartile of the TM-score and RMSD distributions, respectively. The lowest TM-scores of < 0.5 (and highest RMSD values of > 4 Å) were observed for AF2 models of the 8-stranded Opa OMBB (PDB ID 2MLH), which is crucial for the recognition and engulfment of bacterial pathogens *Neisseria gonorrhoeae* or *Neisseria meningitidis* by human cells during pathogenesis [53]. The target structure used is one of the 20 calculated conformers with the lowest energy determined by solution NMR. While the barrel region was predicted accurately, only the extracellular loops did not overlap with the target structure (fig. 2C). This is further highlighted by the superposition-free per-residue Cα-lDDT (fig. S2B), where the scores are higher (> 75) for stranded regions than for the loops (< 50). In this particular case, the flexibility of those loops is in fact essential for the function of the protein, and thus it is not surprising that the AF2 prediction does not match the selected solution NMR structure.

This trend is also observed for other cases where the TM-score is above 0.9, an uncertainty also captured by the predicted Cα-lDDT (pLDDT) computed by AF2. Examples of an 8-stranded and a 12-stranded OMBB are shown in figure 3, highlighting the strikingly good prediction accuracy of pLDDT. Both the Cα-lDDTs and pLDDTs reach values between 95 and 100 for β-strand regions, while the loops (specially those facing the extracellular side of the outer membrane) result in lower lDDT values, with no striking differences between the three AF2 ‘M’, ‘Mnotemp’ and ‘Mreldate’ predictions. Limiting the analysis to the β-strands forming the barrels (which we refer to as ‘barrel cores’ for the remaining text) (fig. 4A), resulted in extremely high median lDDT values of 98.2 ± 1.6, 97.0 ± 1.9 and 97.1 ± 1.9 for the models of the ‘M’, ‘Mnotemp’ and ‘Mreldate’ experiments, respectively; corroborating the marginal deviations observed in the different barrel geometric features.

**Figure 3.**
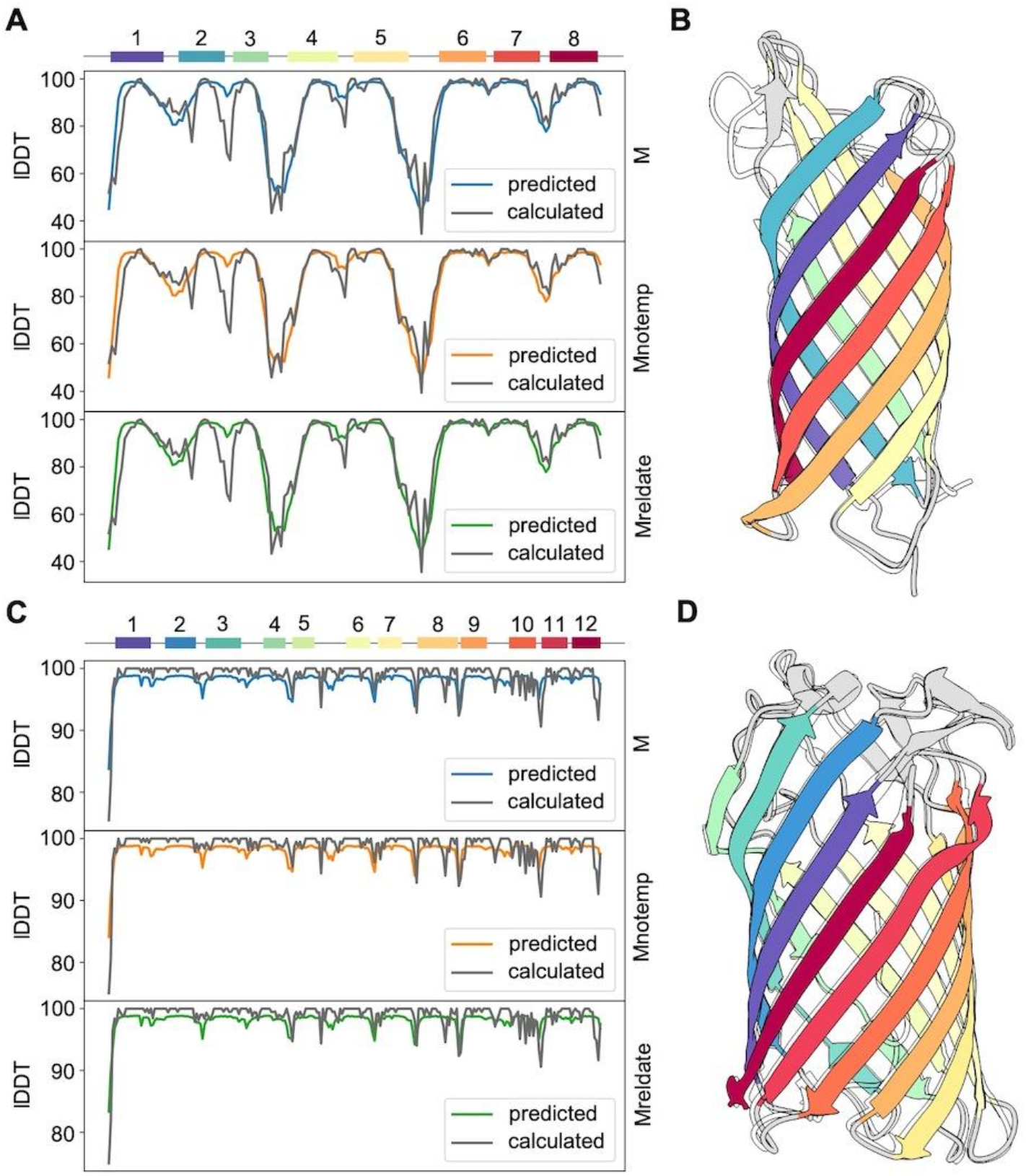
Predicted and calculated Cα-lDDT values per residue for two examples. (A) PDB ID 1P4T, an OMBB with 8 β-strands. (B) and of target PDB ID 4RL8, an OMBB with twelve β-strands. Boxes and numbers indicate the β-strands. The average correlation coefficients between predicted and calculated lDDTs over the three models shown in (A) and (B) are 0.895 ± 0.005 and 0.756 ± 0.009, respectively.

**Figure 4.**
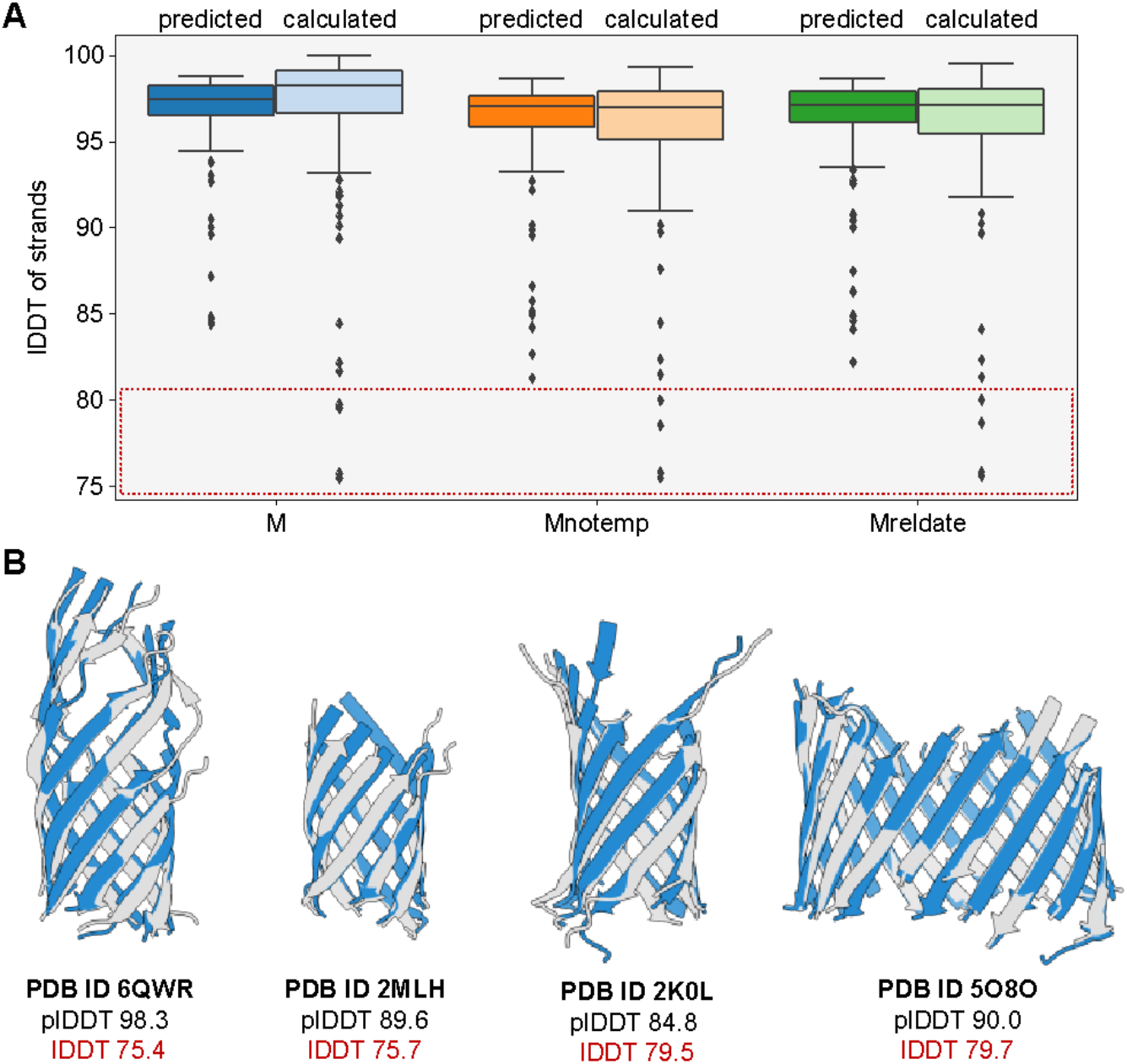
Superposition-free assessment of target structures and AF2 models. (A) Predicted and calculated Cα-lDDT scores of barrel core β-strands are shown in dark and light colors, respectively. (B) The four targets with lDDTs of strands below 80. Shown are only the residues considered as regular regions. Experimental structures and AF2 models are shown in gray and blue, respectively.

Interestingly, while the lDDT and pLDDT correlate well, their distribution for the barrel core regions is different independently of the use of templates (fig. 4A), with AF2 underestimating, on average, their accuracy. In four cases, however, the confidence of the AF2 models for the barrel regions was above 85 while the lDDT was lower, but still within a reasonable range of 75-80 (fig. 4B). The three first cases are PDB IDs 6QWR, 2MLH and 2K0L, all of which are 8-stranded OMBBs whose structures were determined by NMR spectroscopy with 100 or more calculated conformers. The fourth case corresponds to PDB ID 5O8O, the first experimental structure of a 19-stranded mitochondrial import receptor subunit Tom40. Its experimental structure was determined through rigid body docking into a 6.8-Å resolution cryo-EM map of a homology model generated based on the X-ray structure of a homologous mitochondrial voltage-dependent anion channel (VDAC) [54]. While the overall topology of the AF2 model matches this experimental structure and the same residues build up the barrel core, a few strands exhibit a distinct frame-shift along its axis in all three predicted models (fig. S3A), resulting in average lDDT values below 80. A more recent structure of a homologous Tom40 determined by cryo-EM at higher resolution (PDB ID 6UCU) [55], and which was also a target in this study, agrees with the AF2 model (fig. S3B).

Most of these cases, however, were either part of or had full-length homologs in the AF2 training set, thus such high accuracy is expected *a priori.* Of higher interest is the performance of AF2 for cases of novel topology, unknown to AF2. Unfortunately, only one such case is available in the PDB and corresponds to the only known 36-stranded OMBB (PDB ID 6H3I) [44]. It forms the translocon of the Fibrobacteres-Chlorobi-Bacteroidetes type 9 secretion system and its structure was deposited in the PDB after April 30, 2018 (table S1). Although AF2 had never “seen” a 36-stranded barrel, the barrel core and its geometric features were predicted accurately, regardless of the use of templates (fig. 5). In all cases, however, local backbone conformations of the barrel region in the AF2 model are closer to standard geometries of β-sheets than those in the experimental structure. This is not a surprising result as the target is a 3.5 Å cryo-EM structure and lower resolutions lead to higher uncertainties of the atomic coordinates. In the model generated with templates, even the intra- and extracellular loops matched those in the target structure with high accuracy, as also seen in the comparison of predicted and calculated Cα-lDDT values (fig. S2A). There were only minor displacements in the loop regions of the model predicted without any template information.

**Figure 5.**
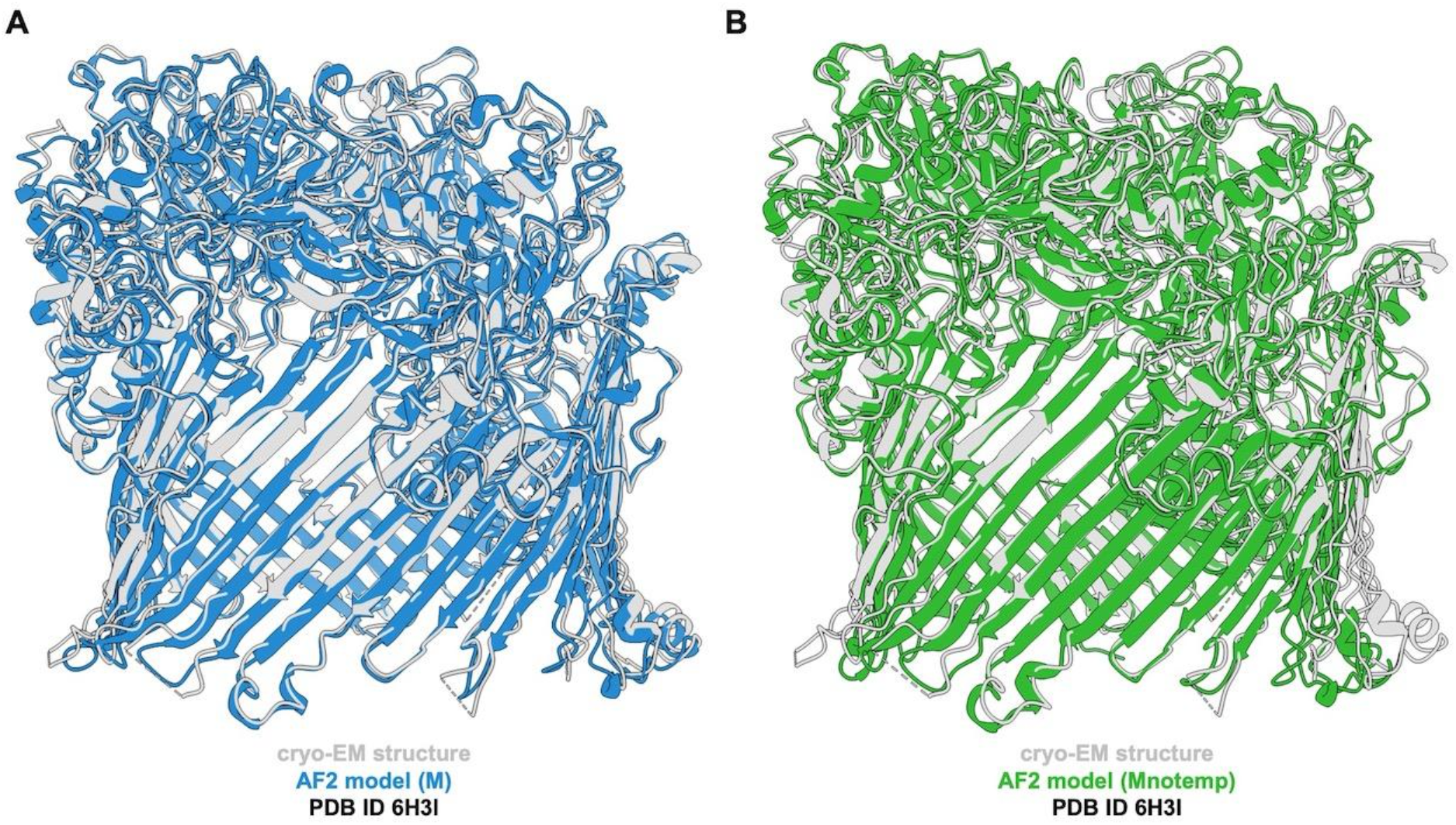
AF2 models of the 36-stranded translocon in the type 9 secretion system (PDB ID 6H3I). Predictions were generated with template (A) and without template (B) information. Excluding templates from the AF2 algorithm resulted in minor displacements in loop regions, while the barrel core was predicted accurately nonetheless. The experimental structure and AF2 models are shown in gray and in color, respectively.

## Discussion

Given the under-representation of transmembrane proteins in the PDB, and consequently in the training set of AF2, it is imperative to evaluate how the algorithm performs for such an important class of proteins. While a study was previously carried out for α-helical transmembrane proteins [14], we focused on the second-largest category: the outer membrane β-barrels (OMBBs), especially those found at the surface of Gram-negative bacteria and their eukaryotic homologs. Gram-positive, multimeric transmembrane β-barrels, which evolved by convergence [56], as well as multimeric OMBBs, which are formed by separate polypeptide/protein chains, such as those forming the anchor domain of trimeric autotransporter adhesins [57], were not considered. We identified 129 non-redundant single-chained OMBBs in the PDB, with topologies ranging from 8 to 36 strands. In all cases, AF2 predictions were highly accurate; all experiments resulted in extremely high median TM-scores above 0.97 and low median RMSD values below 1.4 Å and, overall, no significant differences were observed. For all cases, AF2 correctly predicted the topology of the domain as well as the shear and average diameter, demonstrating that in the case of OMBBs the accuracy of the prediction is not substantially affected by the use or omission of templates.

However, targets with structures deposited in the PDB prior to April 30, 2018 were part of the training set. So even when removing the experimental structure from the template list, structural information might still be used to predict the model as it is stored in the network. The only case in our test set with a topology completely new to the AF2 network was the 36-stranded OMBB from the Fibrobacteres-Chlorobi-Bacteroidetes type 9 secretion system translocon [44]. Although the network has never seen a 36-stranded OMBB, its predictions were highly accurate, even improving on the geometry of the backbone of an experimental low-resolution structure. AF2 predicted correctly the 36-stranded topology, as well as the diameter and the shear of the barrel, but also the intricate folds of the extracellular loops at an extremely high level of detail that translates into an overall lDDT of 86. The models for this test case were of the same accuracy as for those of well-known topologies, indicating that structural information of templates or close homologs is not essential for a correct prediction.

On average, the per-residue lDDT is lower for loops, especially those facing the extracellular side of the outer membrane and independently of the use of templates. This is likely the result of the higher flexibility of extracellular loops observed in experimental structures, which is important for protein function. Such flexibility makes it difficult to predict a static snapshot of those regions at an atomic level of detail, which in turn decreases their pLDDT. Larger differences were also observed for cases where the target was either solved by solution NMR or low resolution cryo-EM. The same was observed in CASP14, where AF2 also performed worst for NMR structures [58]. More recently, Fowler *et al*. examined this by measuring the accuracy of solution NMR structures and comparing them to AF2 predictions [59]. They concluded that, in general, AF2 models are more accurate than NMR ensembles. This is especially the case of β-sheet proteins, which include OMBBs, providing a consistent explanation for the observed low lDDT values when comparing AF2 models with NMR structures. However, when evaluating these values, it must be noted that the lDDT scores are merely a measure of how similar the AF2 models and the experimental structures are, without providing information on which structure is closer to the truth. Still, and although the test set is small, these results provide confidence in the models for OMBBs generated with AF2, especially those with previously unknown topologies.

## Data and code availability

The models generated, as well as the structural features extracted for them and the reference structures, are available in: https://www.modelarchive.org/doi/10.5452/ma-ombbaf2. *barrOs* can be downloaded from: https://git.scicore.unibas.ch/schwede/barrOs. In the repository, detailed instructions on how to use it for the general analysis of proteins with an expected barrel fold are provided, and the HHsearch results used in this work are provided in the Examples folder.

## Acknowledgements

We would like to thank the SWISS-MODEL development team for insightful discussions, technical support and text revisions, and sciCORE at the University of Basel for providing computational resources and system administration support. JP would also like to thank Prof. Andrei Lupas and Jens Bassler (Max Planck Institute for Biology Tübingen) for support in the early days of *barrOs*, and Roger Bamert for contributing to the implementation of the shear number calculation into the pipeline.

## Supplementary Information

**Table S1.**
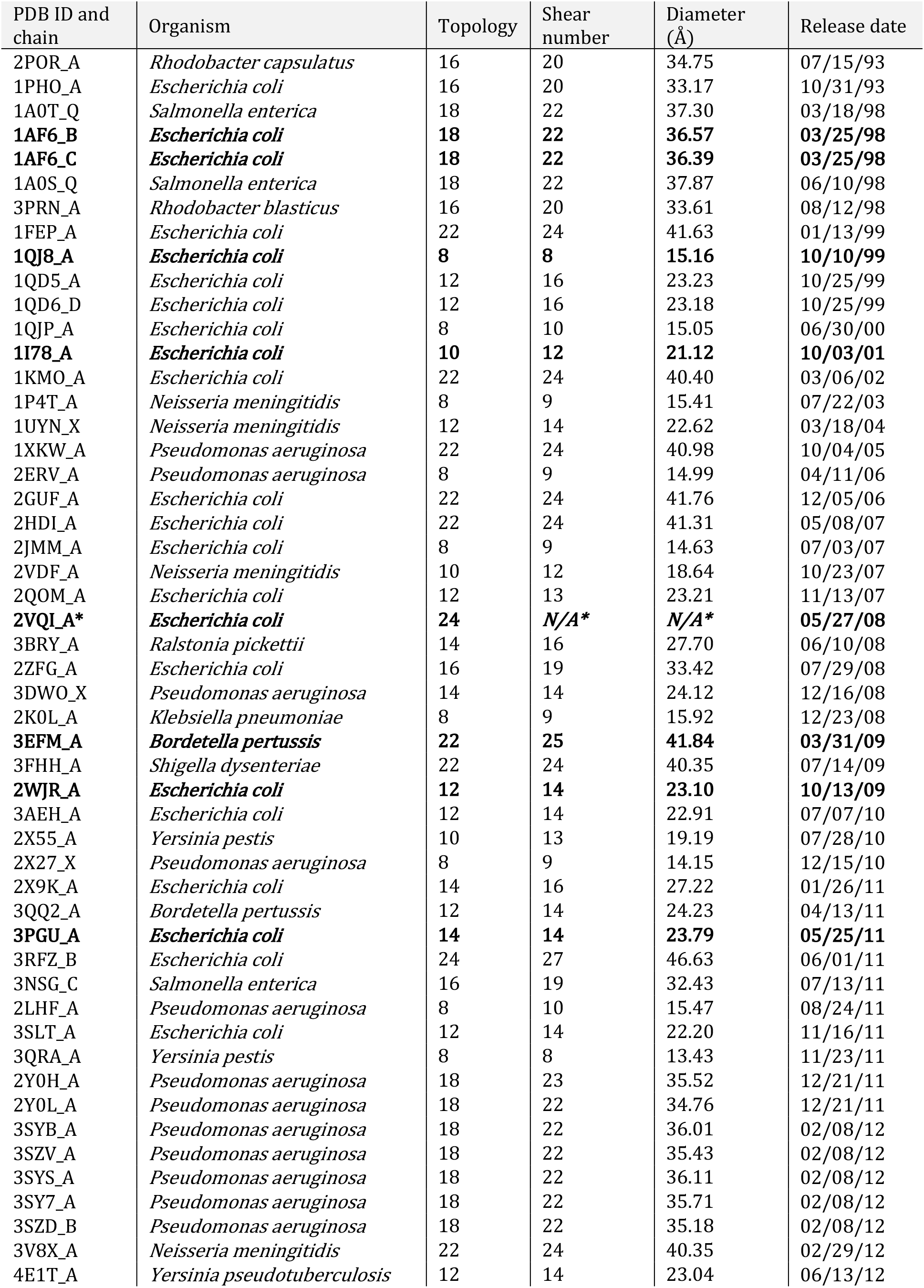
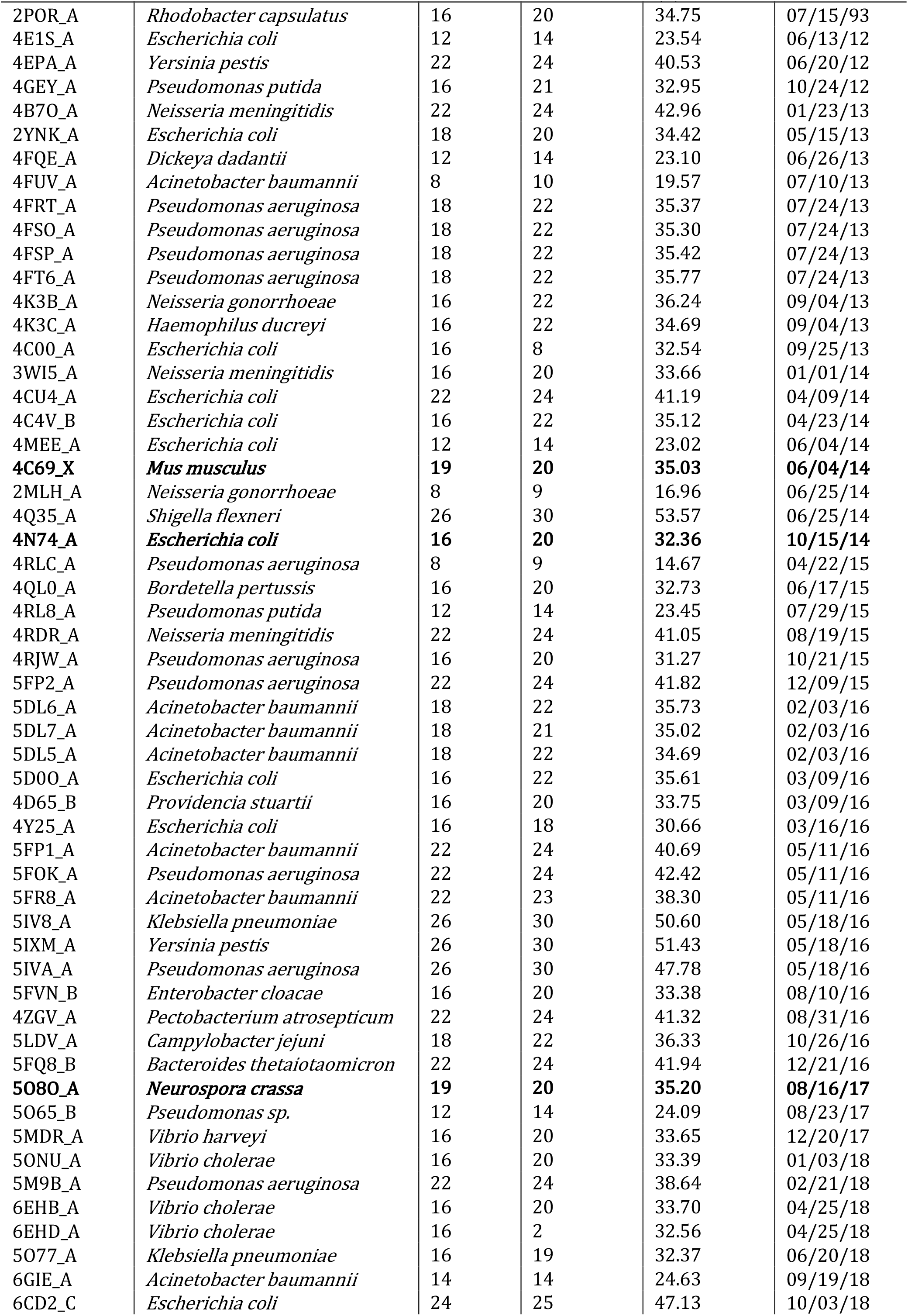
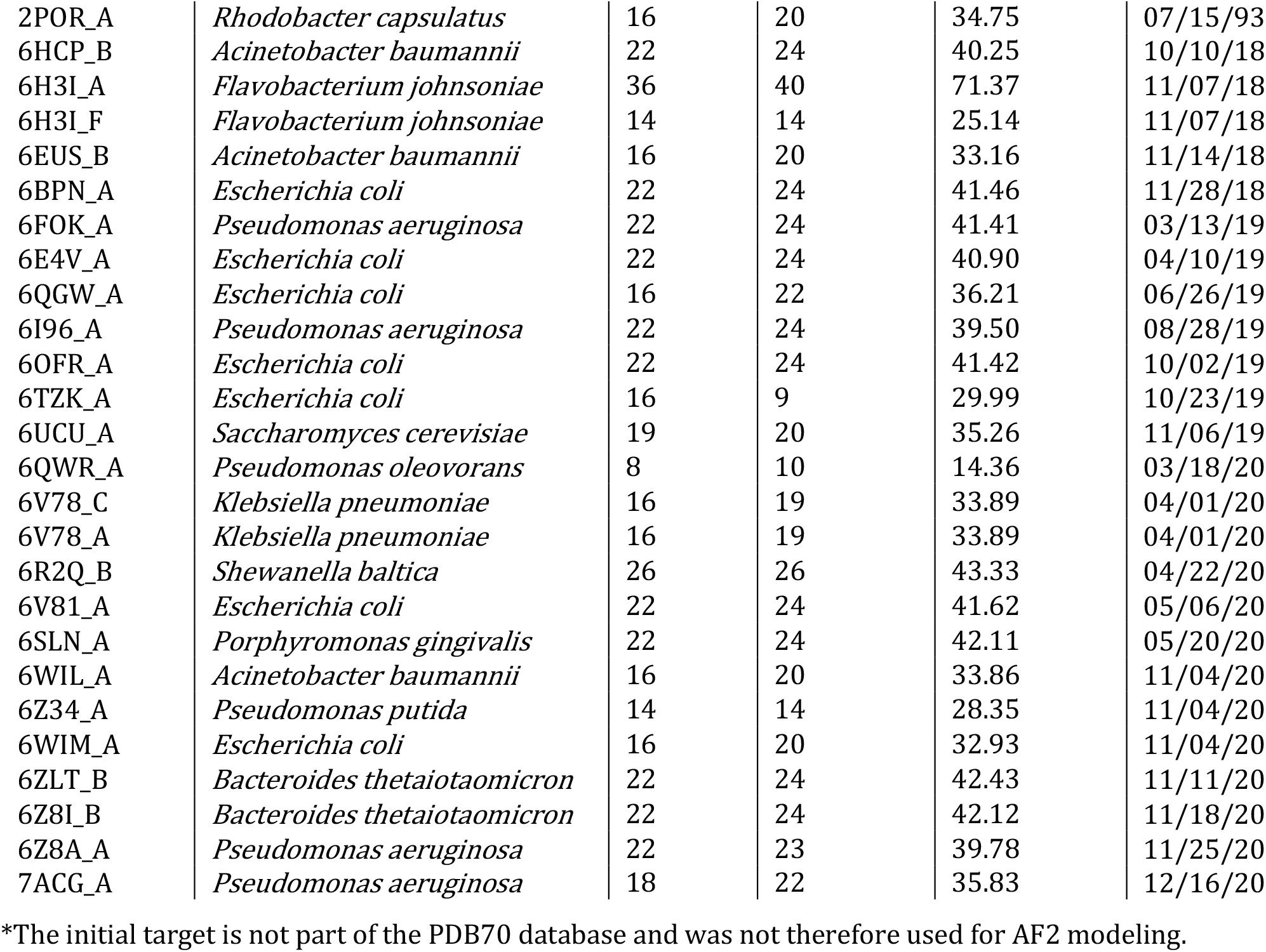
Targets used for AF2 modeling. Initial seeds for HHsearch are shown in bold print.

**Figure S1.**
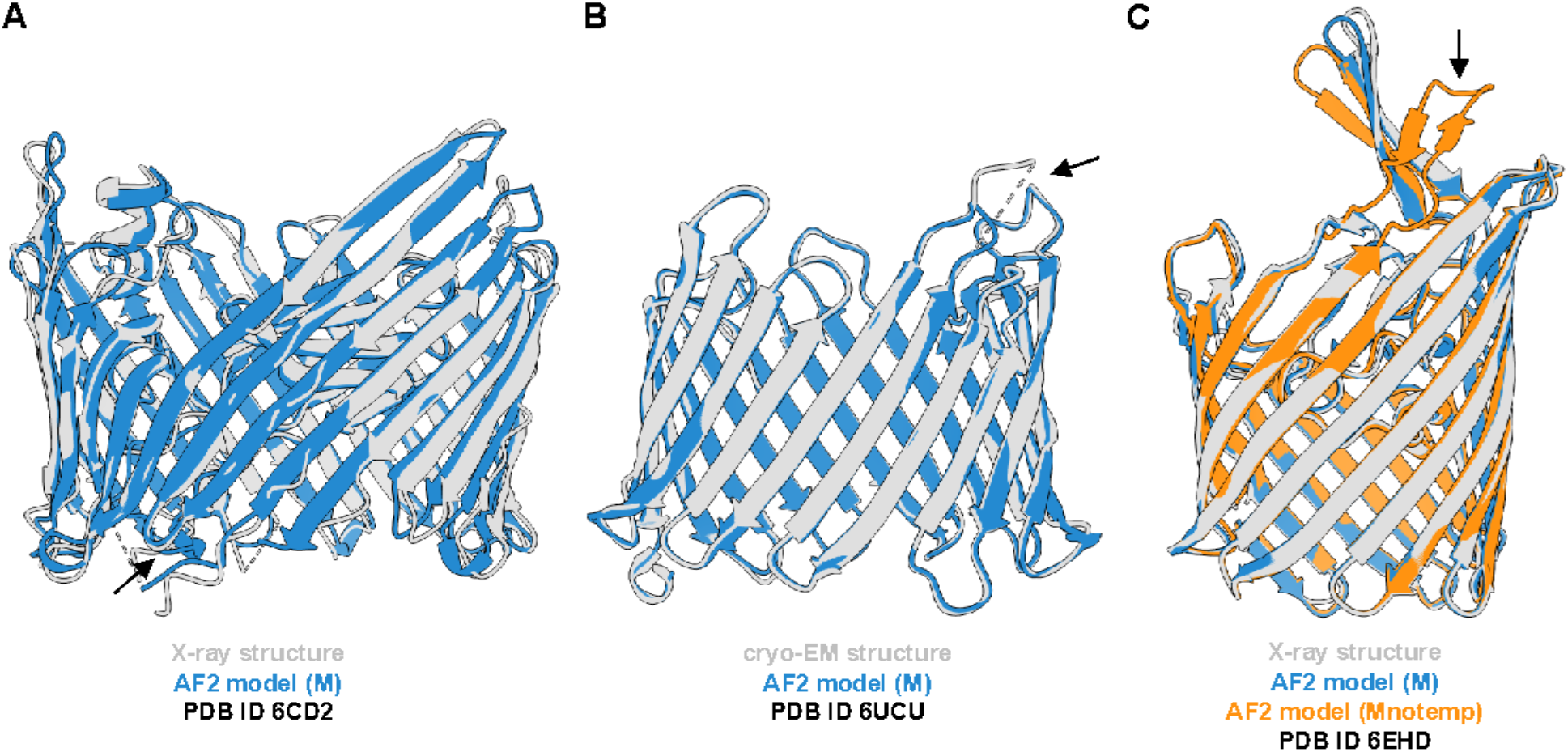
Outliers of the topology analysis. (A) In the AF2 model ‘M’ of target PDB ID 6CD2, two strands were merged into one single regular region due to a short intracellular loop, leading to a miscalculation of a 23-stranded instead of a 24-stranded topology. (B) In the cryo-EM structure of target PDB ID 6UCU, two strands were falsely identified as one due to missing coordinates of an extracellular loop, resulting in a calculated topology of 18 instead of 19 strands. (C) In the AF2 model ‘Mnotemp’, an extracellular loop, which in the experimental structure points inwards into the channel, was predicted facing the outside. Since parts of it were assessed as regular regions by *barrOs*, the calculated shear number differed greatly from those of the X-ray experimental structure. Experimental structures, ‘M’ and ‘Mnotemp’ AF2 models are shown in gray, blue and orange, respectively.

**Figure S2.**
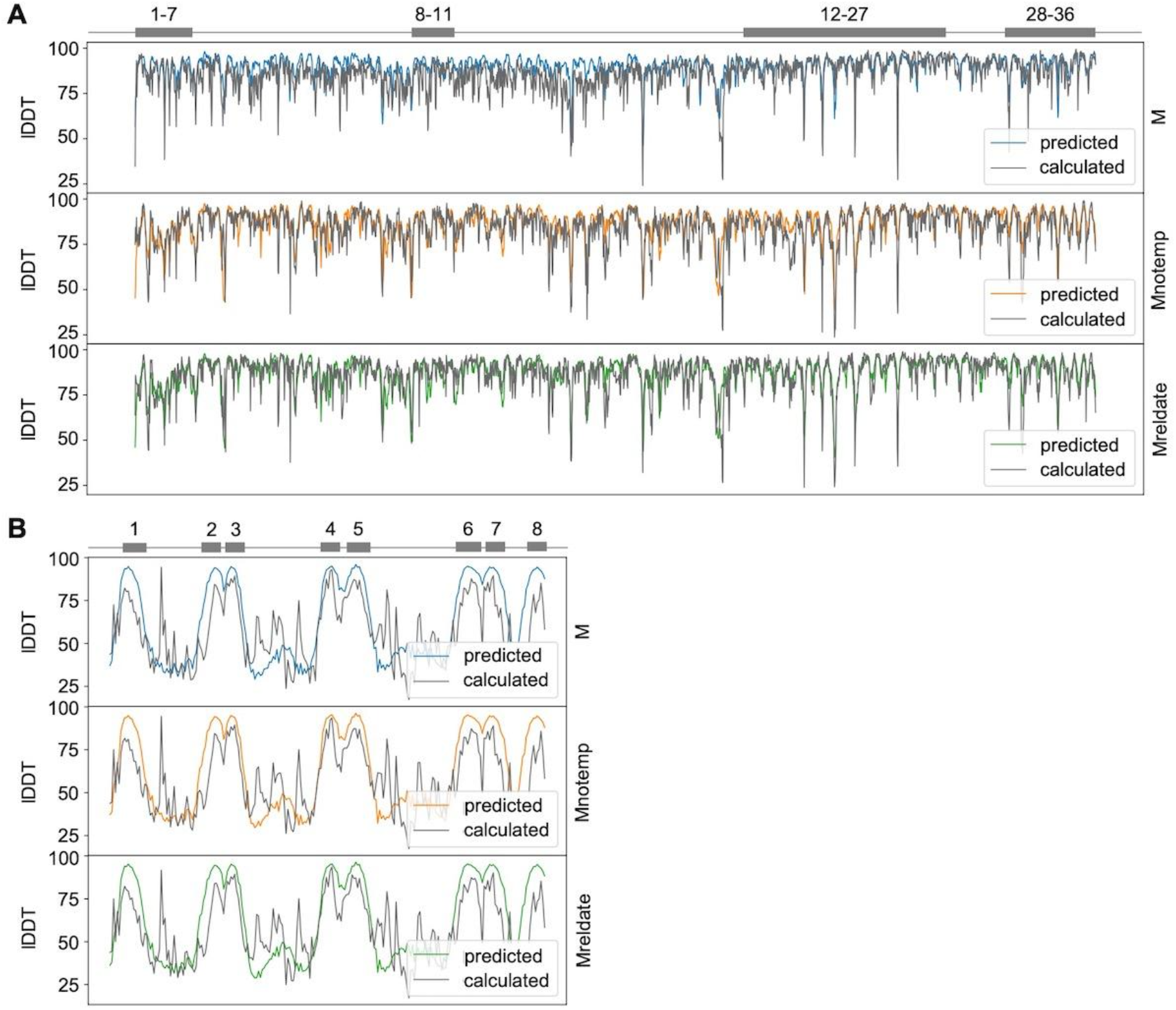
Predicted and calculated Cα-lDDT values for two examples. (A) Target PDB ID 6H3I, an OMBB with 36 β-strands. (B) Target PDB ID 2MLH, an OMBB with 8 β-strands. Gray boxes with numbers indicate β-strands.

**Figure S3.**
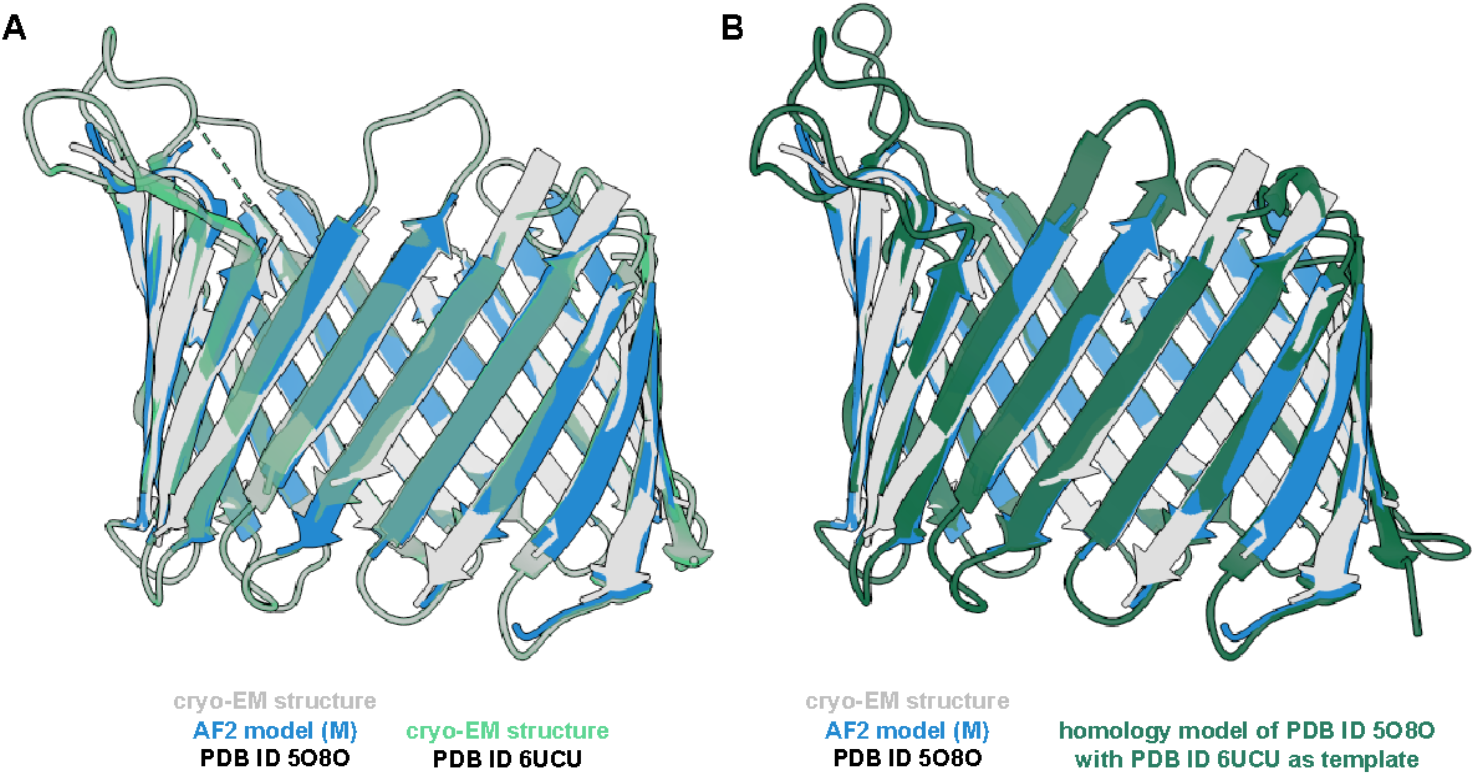
Experimentally determined and predicted structures of Tom40 proteins. The AF2 model (blue) of target PDB ID 5O8O displays a shift in some β-strands as compared to its deposited 6.8-Å cryo-EM structure of *N. crassa* Tom40 (gray). This shift is also observed in the 3.1-Å cryo-EM structure of *S. cerevisiae* Tom40 (PDB ID 6UCU, light green) as well as in a homology model of the target with PDB ID 6UCU as template (dark green). The homology model was generated using SWISS-MODEL [60].

